# *MultiMeta*: an R package for meta-analysing multi-phenotype genome-wide association studies

**DOI:** 10.1101/013920

**Authors:** D. Vuckovic, P. Gasparini, N. Soranzo, V. Iotchkova

**Affiliations:** Department of Medical, Surgical and Health Sciences, University of Trieste, 34100 Trieste, Italy; Medical Genetics, Institute for Maternal and Child Health IRCCS “Burlo Garofolo”, 34100 Trieste, Italy; Human Genetics, Wellcome Trust Sanger Institute, Genome Campus, Hinxton CB10 1HH, UK; Department of Haematology, University of Cambridge, Cambridge CB2 0AH, UK; European Bioinformatics Institute, Wellcome Trust Genome Campus, Hinxton, CB10 1SD, UK

**Author notes:** To whom correspondence should be addressed: Dragana Vuckovic Via dell’Istria 65 34137 Trieste Italy Valentina Iotchkova Genome Campus Hinxton CB10 1HH UK.

## Abstract

**Summary:** As new methods for multivariate analysis of Genome Wide Association Studies (GWAS) become available, it is important to be able to combine results from different cohorts in a meta-analysis. The R package *MultiMeta* provides an implementation of the inverse-variance based method for meta-analysis, generalized to an n-dimensional setting.

**Availability:** The R package *MultiMeta* can be downloaded from CRAN Contact

## INTRODUCTION

Genome-wide association studies (GWAS) have been a powerful tool for genetic discovery for almost a decade. Results have shed light on many different biological processes from lipid metabolism to blood composition as well as social and behavioural patterns (Hindorff et al. 2014). A key to the success of GWAS was the ability to combine several studies in a meta-study, which allowed sufficiently large sample sizes for powered association studies even for variants of small phenotypic effect.

While typically GWAS test one phenotype at a time, many biological features are better described by a combination of several variables. Thus addressing multiple phenotypes can give great increase in power, by taking into account the underlying correlations between variables. Multivariate regression is a straightforward generalization of standard GWAS, where a linear multivariate mixed model can be fit and allows to control for population stratification and relatedness (Korte et al., 2012)(Zhou & Stephens, 2014). In particular, a new set of algorithms included in the GEMMA software (Zhou & Stephens, 2014) allow for a fast multivariate fitting and testing up to ten phenotypes in large sample sizes. As for the univariate GWAS, it might be desirable in some settings to be able to combine multivariate results from different studies in order to further increase statistical power in genotype-phenotype associations.

Here we describe a novel statistically efficient method to perform meta-analysis in a multivariate setting. It is an inverse-variance based method that allows different weights for each cohort in order to take into account the accuracy of each effect estimate. The inverse-variance method has been successfully used for single-trait association testing (Sanna et al. 2008) and is suitable for n-dimensional generalization. Finally it is implemented as part of the R package *MultiMeta* to benefit from flexible environment and open access, as well as extra plotting functions for results visualization.

## METHODS

The multivariate setting implies that results for each single nucleotide polymorphism (SNP) include several effect sizes (or beta-coefficients, one for each trait), as well as related variance and covariance values, since beta coefficients can be correlated.

Let *p* be the number of phenotypes analysed and *n* the number of cohorts to include in the meta-analysis. For each cohort *i* ∈ *{1,…, n}* let *β*^*i*^ be the vector of effect sizes and *Σ*^*i*^ be a *pxp* variance-covariance matrix.

Input data:

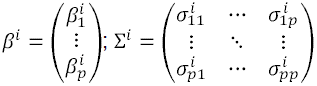

Our method to combine effect sizes is an inverse-variance based method. It is an *n*-dimensional generalization of the single trait meta-analysis, such as the one implemented in METAL software (Willer, Li, & Abecasis, 2010). In particular, each vector *β*^*i*^ is weighted by the inverse of its variance-covariance matrix (Σ^i^)^-1^, then the final effect size *B* is computed as the weighted mean of all the beta coefficients.

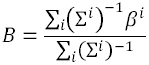

The variance of *B* is:

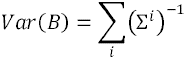

The resulting beta divided by its standard error follows a multivariate normal distribution.

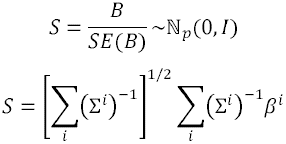

Finally, significance in the multivariate association is estimated from the chisquared statistics with *p* degrees of freedom.

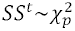

## RESULTS

The method was implemented as part of the R package *MultiMeta*, together with plotting functions useful for visual representation of the results, including Manhattan plot, quantile-quantile plot and an overview plot of effect sizes for a chosen SNP. The default options for the meta-analysis function *multi_meta* are set to work with GEMMA file format (multivariate analysis option). The plotting functions work with output files from the *multi_meta* function by default. However both can be easily adapted to deal with different file formats by changing options, such as field separators, or by changing column names, as specified in the manual. Furthermore the plotting functions can be run by passing files as well as objects in input. This choice is meant to provide more flexibility and avoid unnecessary opening of large files. Example datasets with only few SNPs are included in the package and can be analysed as detailed in the instruction manual. Figure 1 shows two examples of the summary plot obtained with the *betas_plot* function for two SNPs tested on six traits. This is a custom plot, meant to show combined beta coefficients with 95% confidence intervals, as well as correlations between them.

**Figure 1.**
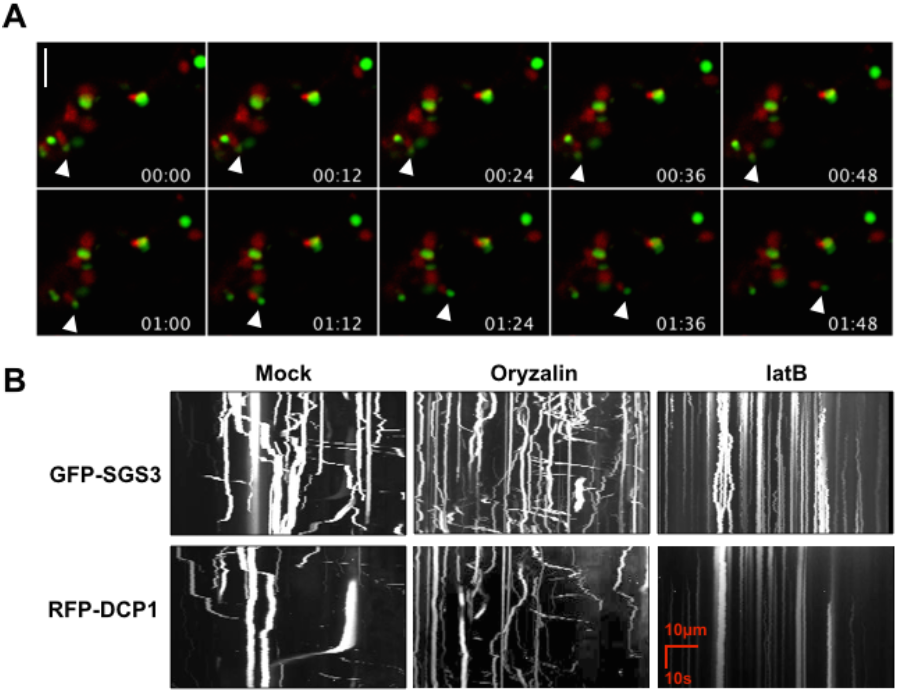
Examples of effect sizes summary plot. This plot provides a visually simple representation of results for a chosen SNP of interest: the left hand panel displays effect sizes with 95% confidence intervals for each trait; the right panel shows a heat-map representing correlations between effect sizes. A) An example of default options in scales of grey; B) For a small number of original cohorts, it is possible to plot effect sizes for each one (light blue and pink on the left panel). It is also possible to choose colours for the heat-map (here: blue, white and pink) and specify the trait names.

The meta-analysis runs on very low memory with default options (e.g. RAM < 250 MB for two cohorts). Computation time is ~0.07 sec/SNP for two cohorts and it grows linearly with the addition of other cohorts in input. As each SNP is analysed singularly, overall computation time depends linearly on the total number of SNPs. By changing settings to increase the dimension of regions in which the genome is divided (see manual), it is possible to increase the performance, while allocating more memory.

The package is freely available on CRAN repository and can be run on any operating system.

## CONCLUSION

Combining results from different cohorts is particularly important for GWAS, where large sample sizes are required to reliably detect alleles with small effects. The R package *MultiMeta* provides a flexible approach to meta-analysing multivariate GWAS and easily visualizing results.

## Acknowledgements

NIcole Soranzo’s research is supported by the Wellcome Trust (Grant Codes WT098051 and WT091310), the EU FP7 (EPIGENESYS Grant Code 257082 and BLUEPRINT Grant Code HEALTH-F5-2011-282510).

## REFERENCES

Hindorff, L. et al. A Catalog of Published Genome-Wide Association Studies. Available at: www.genome.gov/gwastudies. [Accessed August 2014]

Korte, A. et al. (2012) A mixed-model approach for genome-wide association studies of correlated traits in structured populations. Nature Genetics, 44(9), 1066–71.

Sanna, S. et al. (2008) Common variants in the GDF5-UQCC region are associated with variation in human height. Nature Genetics 40:198–203.

Willer, C. J., et al. (2010) METAL: fast and efficient meta-analysis of genomewide association scans. Bioinformatics (Oxford, England), 26(17), 2190–1.

Zhou, X., & Stephens, M. (2014). Efficient multivariate linear mixed model algorithms for genome-wide association studies. Nature Methods, 11(4), 407–9.

